# Rescue Ebola Patients in silico

**DOI:** 10.1101/2020.02.08.939777

**Authors:** Gang Zhao, Sebastian Binder, Georg Pongratz, Michael Meyer-Hermann

## Abstract

Ebola, one of the most pathogenic and lethal viruses, repeatedly leads to severe outbreaks with large numbers of casualties. By integration of data of fatal, non-fatal and asymptomatic cases, we developed a mathematical model of the immune response to Ebola infections and found that TNF-α can explain subsequent cytokine responses and allows to quantitatively stratify patients into survivors and fatal cases. The model further suggests that antibodies provide the key resolving mechanism in survivors while IFN-γ is the key defense mechanism in fatal cases. Exogenous IFN-γ and inhibition of excessive inflammation have the potential to rescue fatal Ebola infections in silico. In view of the high mortality of this disease, limited real life studies, and the unforeseeable next outbreak, our in silico analysis is meant as a guide for future research by the determination of the most efficient strategies for Ebola treatment.

---

Ebola virus (EBOV) is a member of the *Filoviridae* family, a group of viruses highly pathogenic in humans and nonhuman primates. EBOV is subdivided into several species, with Zaire being the most lethal one. The case fatality rate of Zaire EBOV is 60%-90%, and death usually occurs within 5-7 days after symptom onset^1, 2^. Fatal cases of Ebola virus disease (EVD) are characterized by uncontrolled virus replication, excessive release of inflammatory mediators by the host, lymphocyte apoptosis, coagulation abnormalities and organ damage^3–5^. The requirement of biosafety level 4 containment for the study of live virus strongly limits available data.

*In vitro* studies suggest that monocytes/macrophages and dendritic cells respond differently to EBOV infection. Monocytes/macrophages secrete a large amount of cytokines and chemokines upon infection *in vitro*. This is largely independent of virus replication as UV-treated virus particles elicited similar transcriptional activation and cytokine/chemokine secretion^6^. However, type 1 interferon production and response are inhibited by various EBOV proteins^7–9^. EBOV grows exponentially in primary human dendritic cells (DC); however, DC fail to secrete pro-inflammatory cytokines and their T cell stimulatory capacity is inhibited^10, 11^. These results are in line with the notion that EBOV is able to cause global immunosuppression. Despite significant lymphocyte apoptosis, there is a rebound of the number of lymphocytes in a late-stage of EBOV infections in nonhuman primates^12^, mice^13, 14^, and humans^15^ suggesting the activation of the adaptive immune response. Adoptive transfer studies showed that whole splenocytes from moribund EBOV infected mice protect naïve animals from EBOV challenge^13^. Specific antibody responses have also been observed in some EBOV and other filovirus infected fatal human cases^16, 17^. These results suggest that although insufficient for control of the virus in the late phase, specific CD8+ T cell and B cell responses can be generated in fatal cases.

Patients surviving EBOV infection are characterized by early appearance of a pro-inflammatory response and an EBOV specific humoral response in blood, and EBOV specific cellular responses concomitant with virus clearance^16, 18, 19^. Fatal cases show late and uncontrolled pro-inflammatory cytokines in blood until death, with parallel very high levels of anti-inflammatory cytokines, which lead to a “cytokine storm”^5, 20^ that is directly related to disease pathology^12, 21, 22^. EBOV infected asymptomatic humans are characterized by extraordinarily strong and rapidly resolved inflammatory responses followed by transient cytotoxic T cell activation and moderate IgG and IgM response^3, 23^.

Taken together, current information supports the notion that the timing and the kinetics of viral replication and of the immune response is key to the outcome of infection and that by accelerating the immune response or retarding viral replication, the chance of survival might be substantially increased.

In view of limited experimental possibilities, mathematical models are important tools in understanding the race between viral replication and anti-viral immune responses. If validated by human patient data, such models could be used to stratify patients and to propose optimal treatment strategies. So far, only few mathematical modeling studies of EBOV infection exist in literature^24, 25^; however, none of them aimed at explaining an *in vivo* human data set.

In this article, we developed a mathematical model for the cytokine response in EBOV infection, and parameterized the model by integration of different sets of human patient data for survivors, fatal cases^18, 20^ and asymptomatic infections^3, 23^. We tested different hypotheses for the reason of fate decisions when fitting the model to the data sets. The results suggested that a differential early inflammatory immune response to the virus can explain the whole history of the immune response of survivors, fatalities, and asymptomatic infection, while differential properties of the virus, or differential properties of the adaptive immune response, cannot explain the data. This model allows for a stratification of EBOV patients into survivors and fatal cases at the time of presentation in the clinics. Moreover, the model also suggested that fatal cases could be rescued by increasing and accelerating an effective T cell response and inhibiting excessive inflammation in a fine-tuned manner.

## Result

### A model of cytokine response to Ebola infeciton

A cytokine response model (Fig. 1) was developed and tailored to the availability of data on infection and immune response (see supplement for the details of the modelling approach). Here, we briefly introduce parameters that are important for understanding the following sections. The virus replicates with a rate *p* and is removed by innate (represented by TNF-α), by cellular (represented by IFN-γ), and by humoral immune responses (represented by antibodies) with a rate of *R*_f_, *R*_g_ and *R*_a_ respectively. TNF-α induction by virus and IFN-γ is described by *Q*_vf_ and *Q*_gf_ respectively. IFN-γ induction by virus, TNF-α and itself is described by *Q*_vg_, *Q*_fg_, and *Q*_gg_, respectively.

**Fig. 1.**
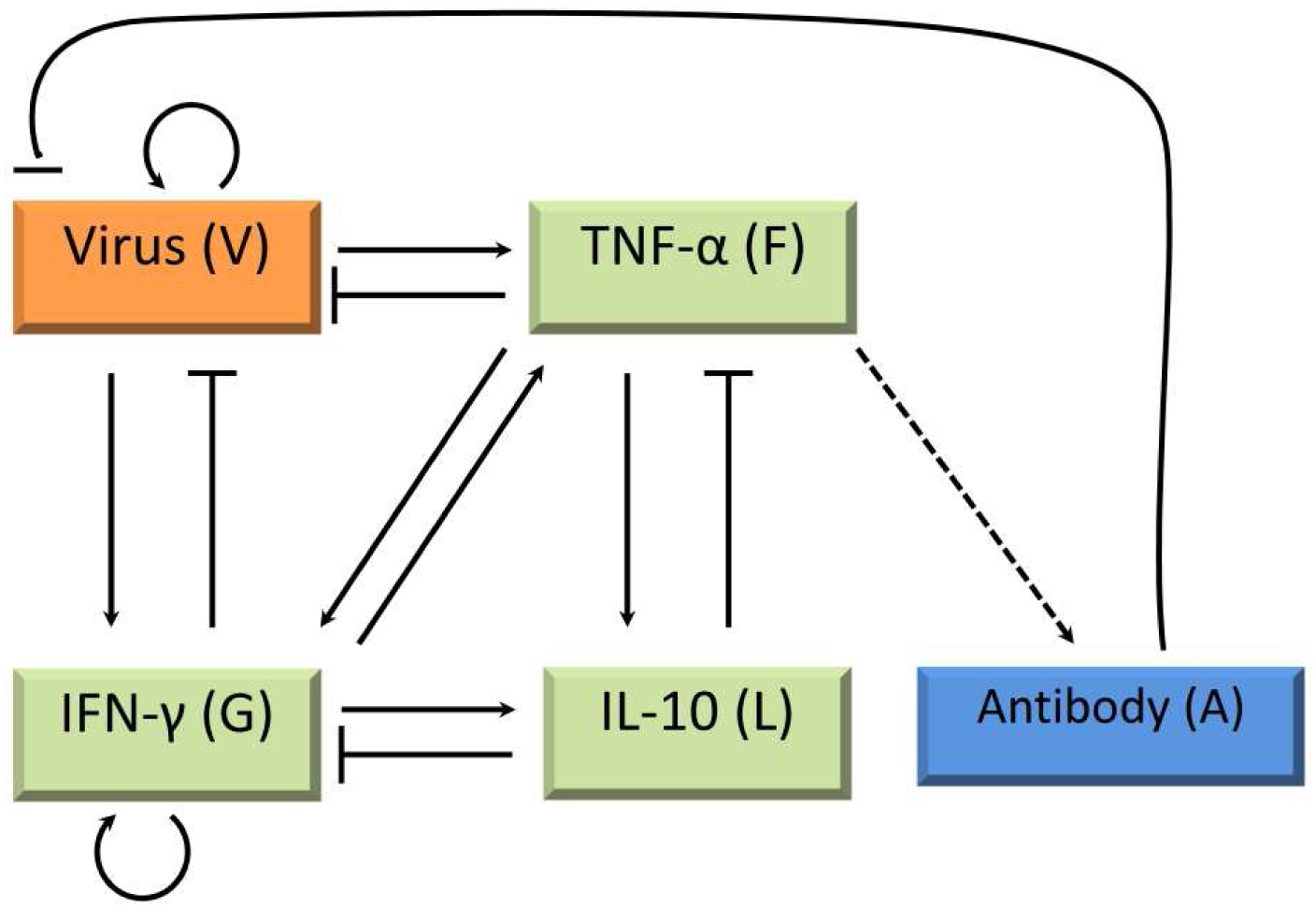
Scheme of the cytokine network model. Arrows represent induction, lines with bar represent inhibition. The broken arrow (from TNF-α to antibody) stands for a delay of antibody induction.

### The fate of the EBOV patient is derived from differential TNF-α responses

With the aim to explore potential differences between EBOV fatalities, survivors, and asymptomatic infections in the inoculation phase, we carried out numerical optimization studies to test the following hypotheses:

H1) Initial viral dose and virus replication rate (*p*) are different among the three groups;
H2) Virus clearance rate by the innate and the adaptive immune response (*R*_f_ and *R*_g_) are different among the three groups;
H3) The innate immune responses (*Q*_vf_ and *Q*_gf_) are different among the three groups;
H4) The adaptive immune responses (*Q*_vg_, *Q*_fg_ and *Q*_gg_) are different among the three groups.
H5) Both the innate and the adaptive immune responses (*Q*_vf_ and *Q*_vg_) are different among the three groups.

H1, H2 and H4 failed in explaining the data while H3 and H5 were compatible with the data. The best fit was obtained from H3 (Fig. 2, see also Fig. S1 for all the accepted fitting results from H3). The relative likelihood of the best result from H5 (Fig. S2, see also Fig. S3 for all the accepted fitting results) is 0.69, meaning that H5 is 0.69 times as probable as H3 to simultaneously explain the three data sets. To make sure that the specific choice of the differential parameters does not influence the result, we also tested additional variations of the hypotheses (see Supplement), but none was compatible with the three data sets.

**Fig. 2.**
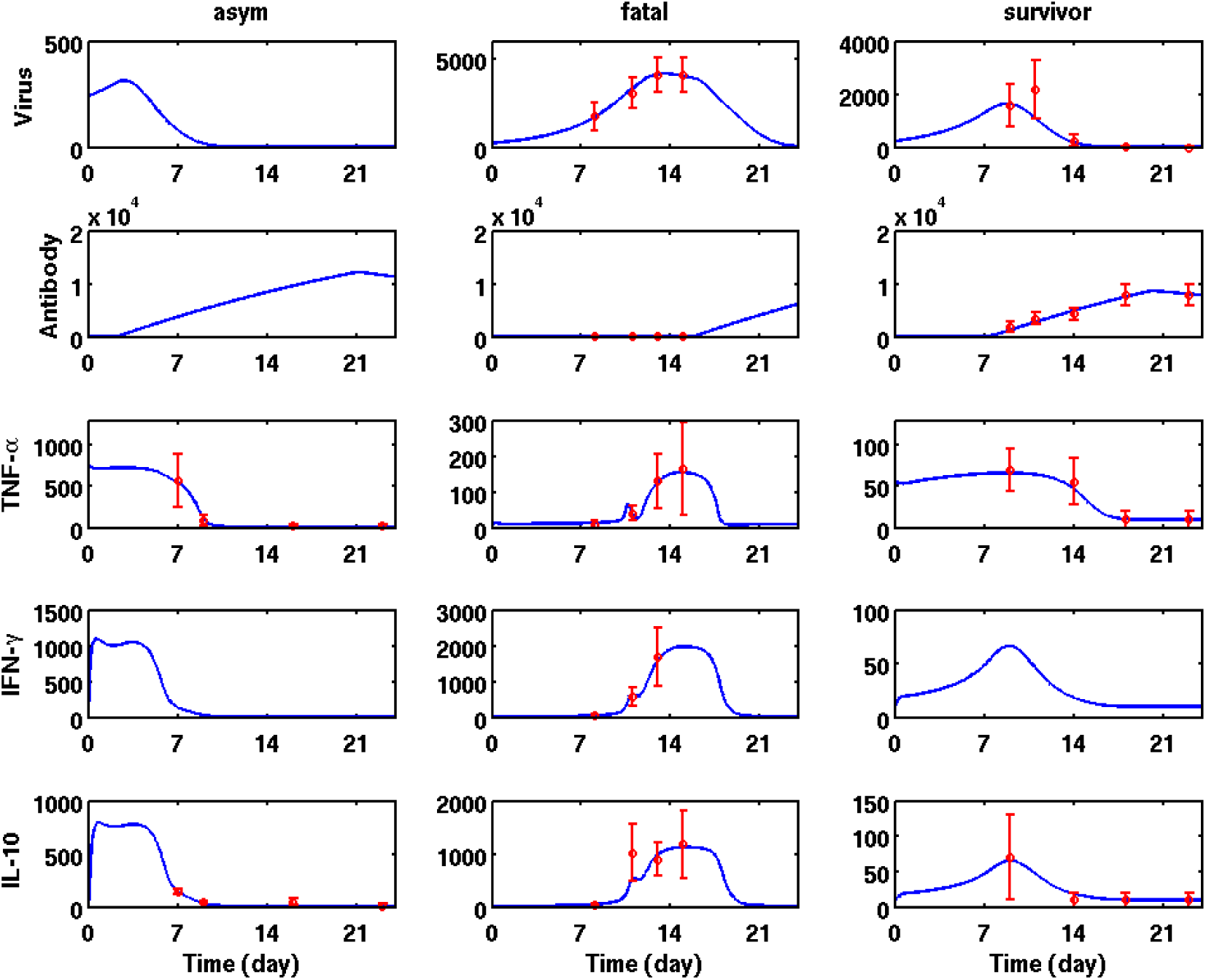
Differential innate immune response (H3) explains best the data of the three groups simultaneously. Each column is corresponding to one group. Note that the fatalities succumbed on day 16 or 17. Circles with error-bar are measurements, curves are the simulation result.

In order to find a fate decisive parameter, we analyzed the distributions of parameters in H3 (Fig. 3A and B), and H5 (Fig. 3C). For both models (H3 and H5), *Q*_vf_, which represents TNF-α induction by EBOV, showed a sequence consistent with disease severity, with the asymptomatic infections being most and the fatalities being least sensitive to EBOV. In contrast, *Q*_gf_, which represents TNF-α induction by IFN-γ, of the three patient groups were comparable (in H3, Fig. 3A), and *a*_F_ (in H3, Fig. 3B), which represents the maximal TNF-α production rate, was smallest in survivors and largest in asymptomatic infections. *Q*_vg_, which represents IFN-γ induction by EBOV, showed a similar pattern as *a*_F_ (in H5, Fig. 3C). The values of other parameters are shown in the supplement Table 1 for both H3 and H5. We conclude that TNF-α induction by EBOV has the potential to be a fate decisive parameter.

**Fig. 3.**
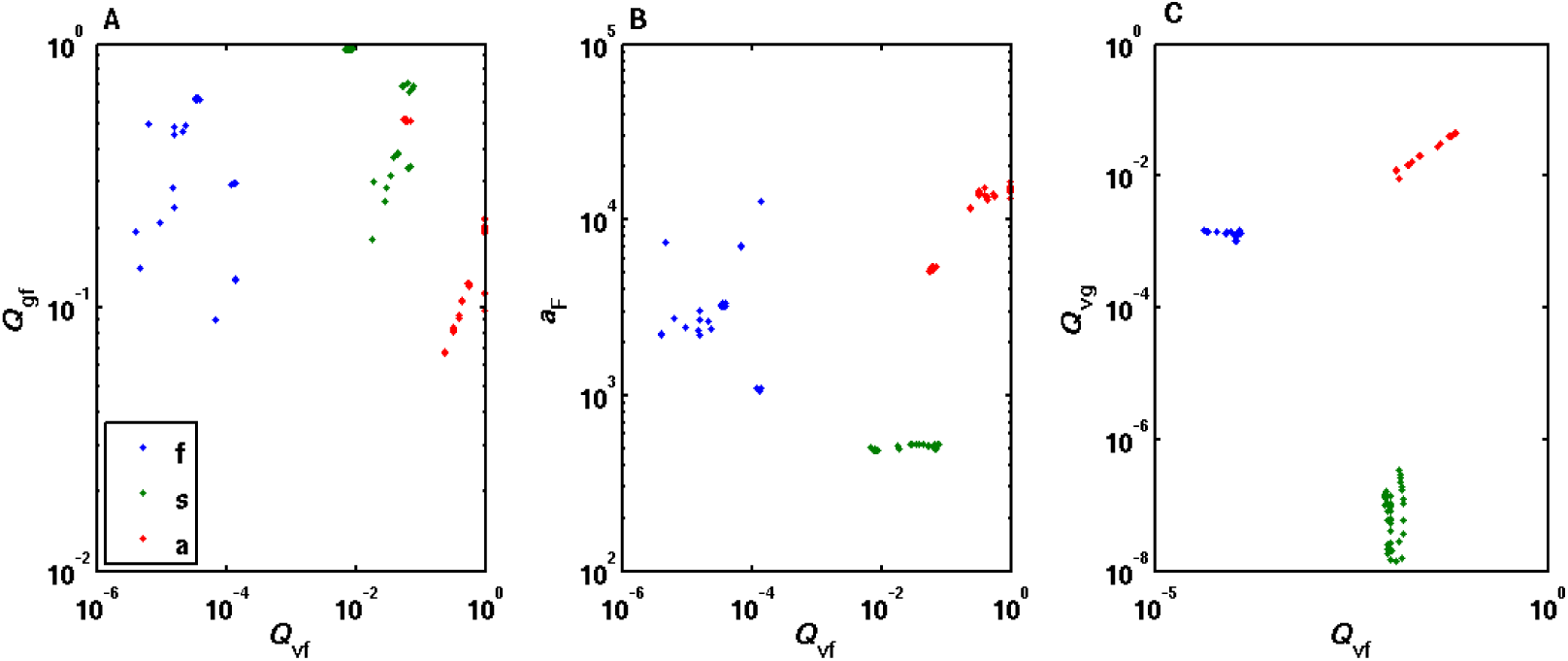
Differential parameters of the three groups (f, fatalities; s, survivors; a, asymptomatic infections) from all the accepted fitting results (A and B for H3, C for H5). Only *Q*_vf_, which represents TNF-α induction by EBOV, showed a sequence consistent with disease severity in both H3 and H5. Please note that *a*_F_ is not a free parameter during the optimization procedures for H3. It is influenced indirectly by *Q*_gf_ in the steady state constraint of TNF-α..

**Table 1.**
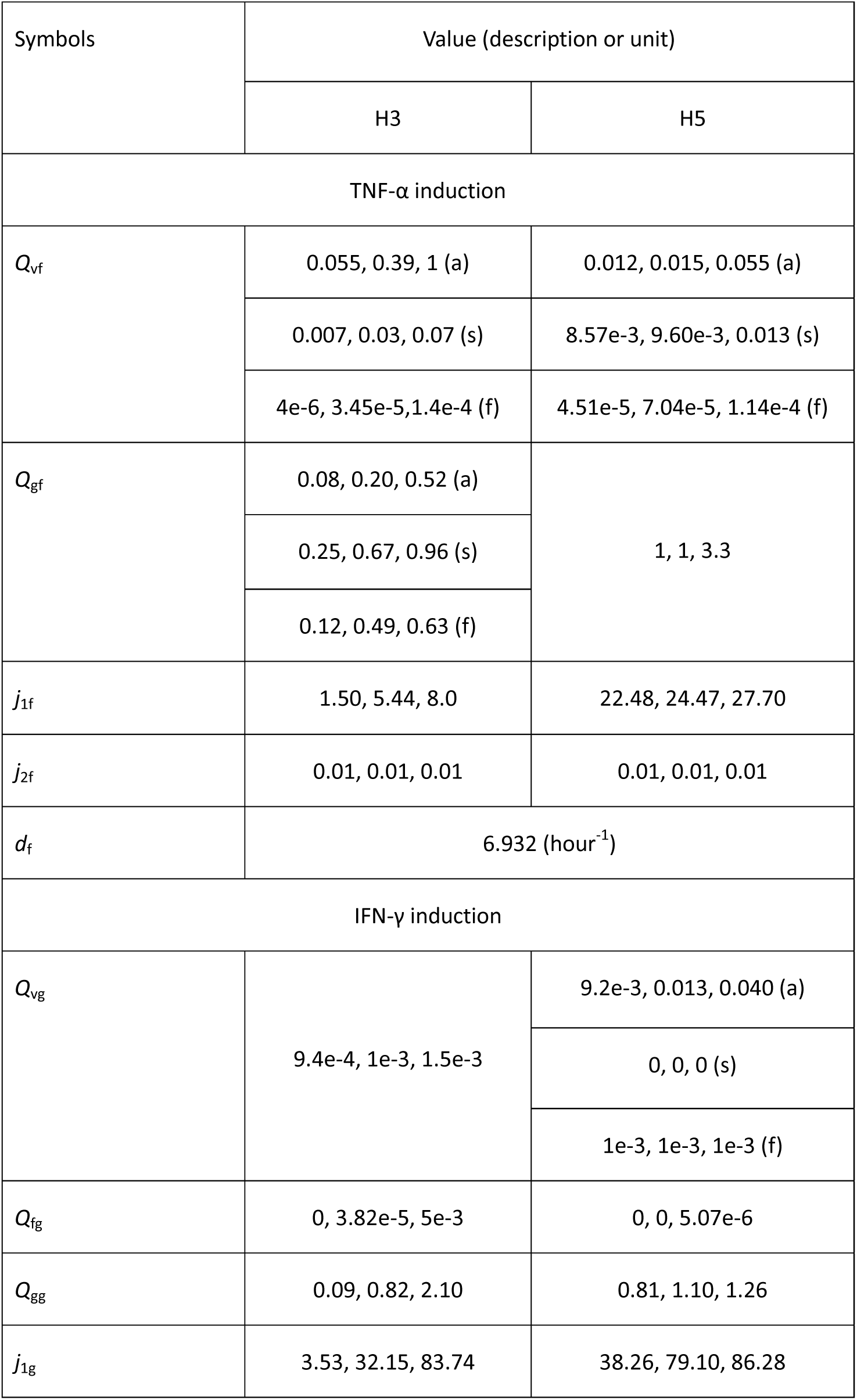
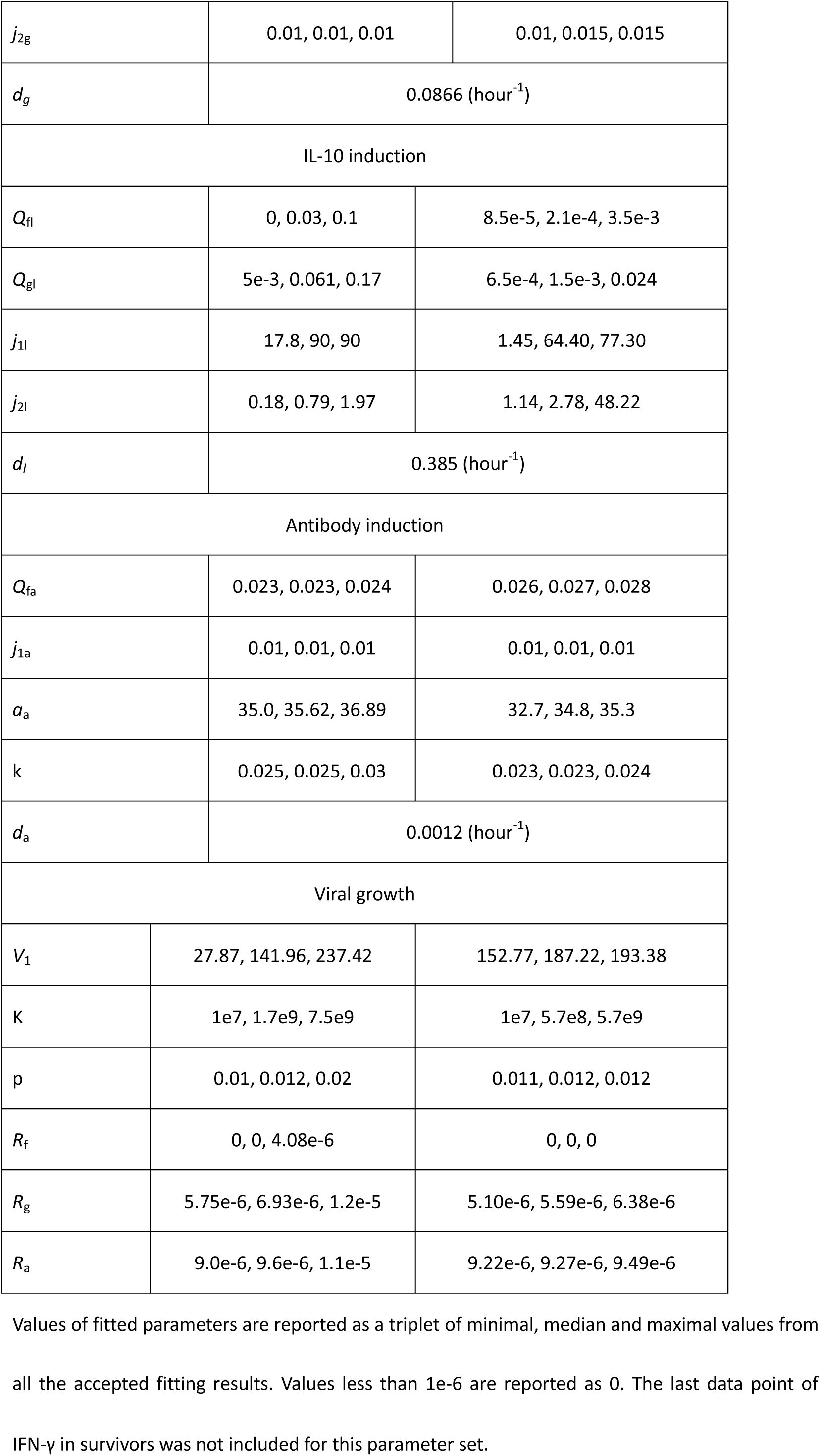
Parameter values for hypotheses H3 and H5

Since TNF-α response has been identified as the earliest difference that determined the final fate, we then asked whether a single measurement of TNF-α in the early phase could be used to predict the final outcome. We constructed five virtual patients whose TNF-α level on day 8 (the day of clinical admission) are 20, 30, 40, 50 and 60 pg/ml, respectively. The mean TNF-α level of survivors and fatal cases on day 8 are 70 and 10 pg/ml, respectively. We fitted the virtual patients, who have only one data point, together with the other three groups using hypothesis H3 (differential innate immune responses). The results suggested that a virtual patient mimics the fatalities, as quantified by the area under curve (AUC) of the viral load, if the TNF-α level on day 8 is 40 pg/ml or less and that a virtual patient could survive EBOV infection if his TNF-α on day 8 is 50 pg/ml or more (Fig. 4).

**Fig. 4.**
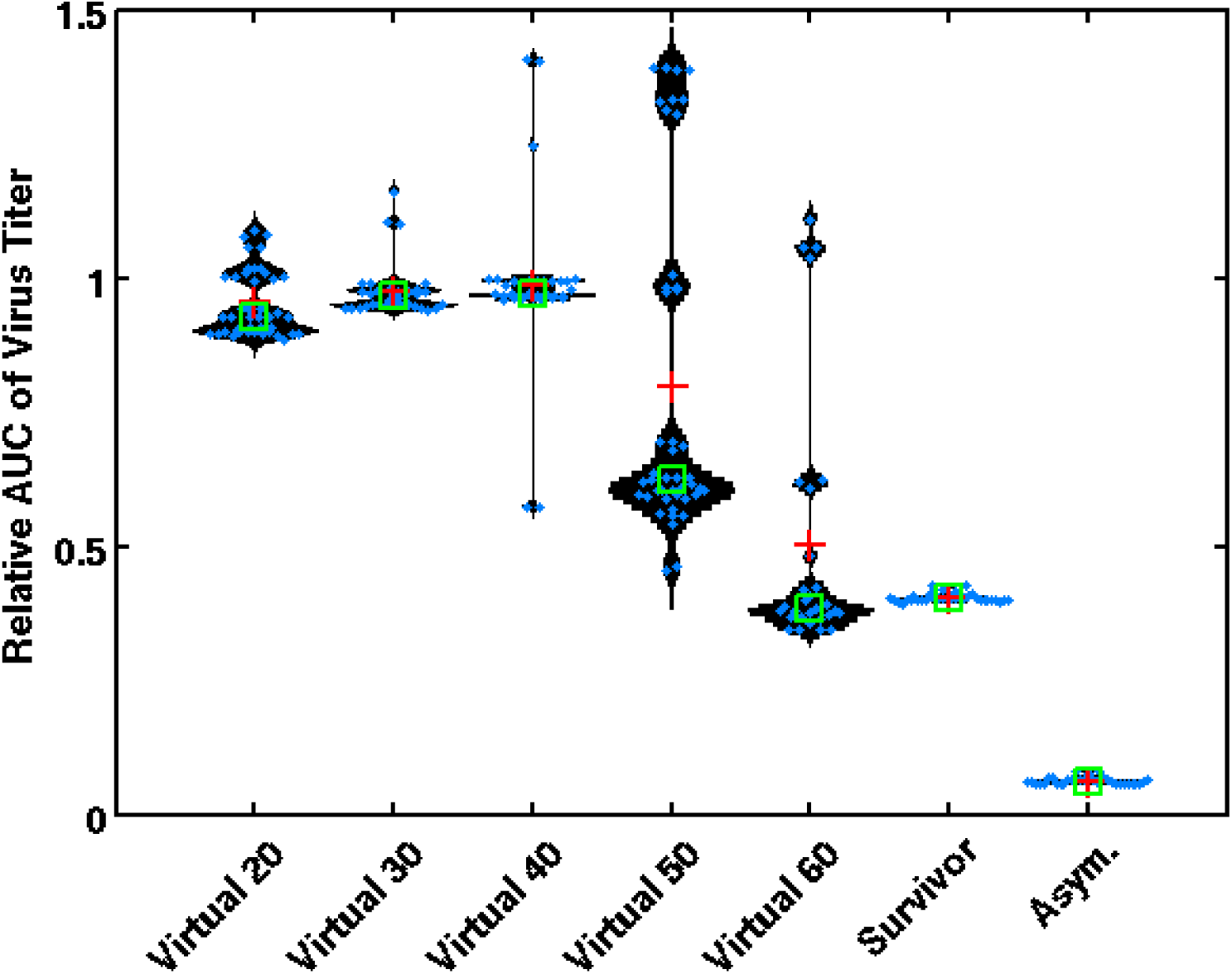
Stratification of virtual patients based on early TNF-α levels. The viral load of virtual patients, survivors and asymptomatic infections, normalized by that of fatal cases are shown. Blue points are results from 100 independent numerical optimizations. Green squares and the red crosses denote the mean and the median value, respectively. “Virtual X” means the virtual patient whose TNF level is X pg/ml on day 8 after infection. AUC is calculated from day 0 to 15 (one day before fatal cases succumbed to the disease.)

### Early immune response to EBOV suggested by the model

The cytokine response after EBOV infection, as indicated by fitting results from both H3 and H5 (Fig. 2, S1 and S2), can be summarized as follows. For survivors, the TNF-α response was initiated early after infection. The level of TNF-α reached five times of its resting level between day 0 and 4. IFN-γ either peaked on day 8 to more than five times the resting level (H3, Fig. S1) or was downregulated after infection (H3 and H5, Fig. S1 and S3). Interestingly, although we did not include IFN-γ levels of survivors in the numerical study, a subset of the fitting results reproduced the reported values (< 20 pg/ml in^18^). However, we kept those fitting results that showed IFN-γ upregulation in our analysis since IFN-γ mRNA in circulating blood mononuclear cells from survivors showed such a trend^18^. Antibody levels exceeded a titer of 100 on day 7 and increased continuously afterwards.

For fatal cases, IFN-γ reached five times its resting level on day 7. TNF-α did this on day 11. The antibody exceeded a titer of 100 on day 15 only, 1-2 days before death of the patients.

For asymptomatic cases, TNF-α reached fifty times the resting level on day 3 at the latest. IFN-γ is either downregulated below resting level (H3), or upregulated to around 100 (H5) and 1000 (H3) pg/ml. Antibody levels exceeded a titer of 100 around day 3.

Viral growth rate and viral clearance rates induced by different immune factors did not differ between the parameter sets of H3 (differential innate immune response) and H5 (differential innate and adaptive immune response) (Fig. 5), which supports the credibility of the results.

**Fig. 5.**
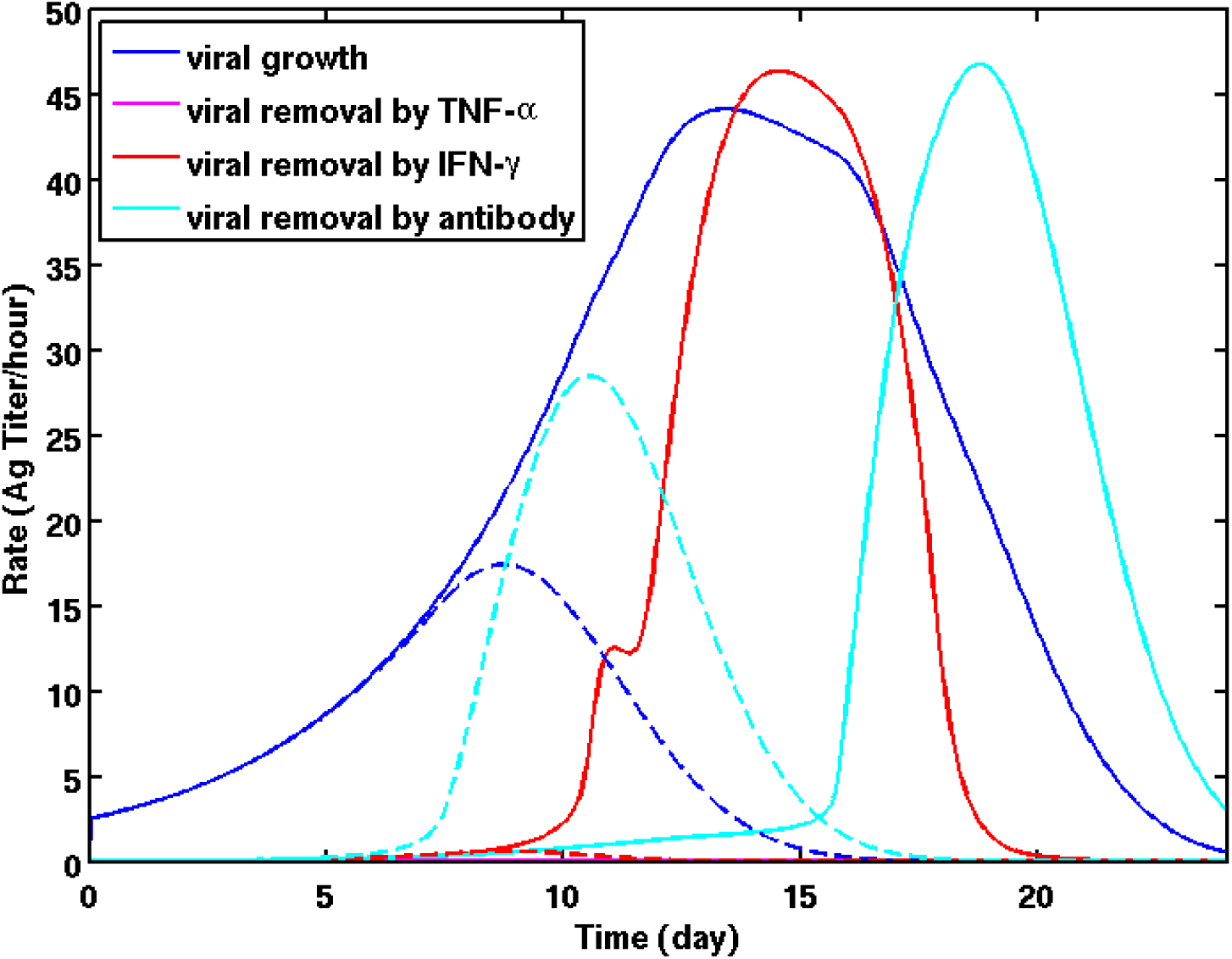
Natural history of EBOV growth rate (blue), and rates of removal by immune factors (see figure inset) in the fatalities (full line) and survivors (broken line). TNF-α induced viral clearance is negligible in both groups of patient. Although insufficient, IFN-γ was the main defense arm against EBOV in the fatalities, while antibody removed most of the viruses in the survivors. The fatalities succumbed on day 16 or 17. These curves were based on the best fit in Fig. 2.

Interestingly, there was no difference between survivors and fatal cases before day 7 (Fig. 5). TNF-α induced viral clearance is negligible in both groups all along. For survivors (broken lines in Fig. 5), antibody-induced viral clearance was the main resolving mechanism, which appeared on day 7 and surpassed the viral growth rate on day 9. For the fatalities (full lines in Fig. 4), IFN-γ-induced viral clearance increased from day 9 and constituted the main defense mechanism, although antibody also contributed a small part after day 13, when the viral growth rate started to decrease. Although IFN-γ is suggested to have nearly the same capacity in virus clearance as antibody (5.76e-6 vs 9.06e-6, and see the possible ranges in Table 1), IFN-γ rose in the fatalities 2 days later than antibody did in the survivors (Fig. 5). We conclude that survivors and fatal cases differ in the timing of the immune response onset and in the choice of the dominant immune strategy.

### Immune-interfering therapy for Ebola patients

Considering that both high viral load and excessive inflammations in fatal cases contribute to the multiple pathologies that finally led to death, we carried out simulations to explore the possibility of limiting viral replication and inflammation at the same time, by modulation of multiple aspects of the immune response. Fig. 6 shows the results of one example, in which exogenous IL-10 and IFN-γ were added to fatal patients, with fixed doses, every 8 hours from day 8, i.e. the first day after clinical admission. The simulation results suggested that viral load and inflammation are better controlled than in the untreated fatal case and that the virus is cleared without excessive inflammation. A scan of the dose-space of IL-10 and IFN-γ suggested that a successful rescue can only be achieved if the doses of IL-10 and IFN-γ are confined to a small region (Fig. S4).

**Fig. 6.**
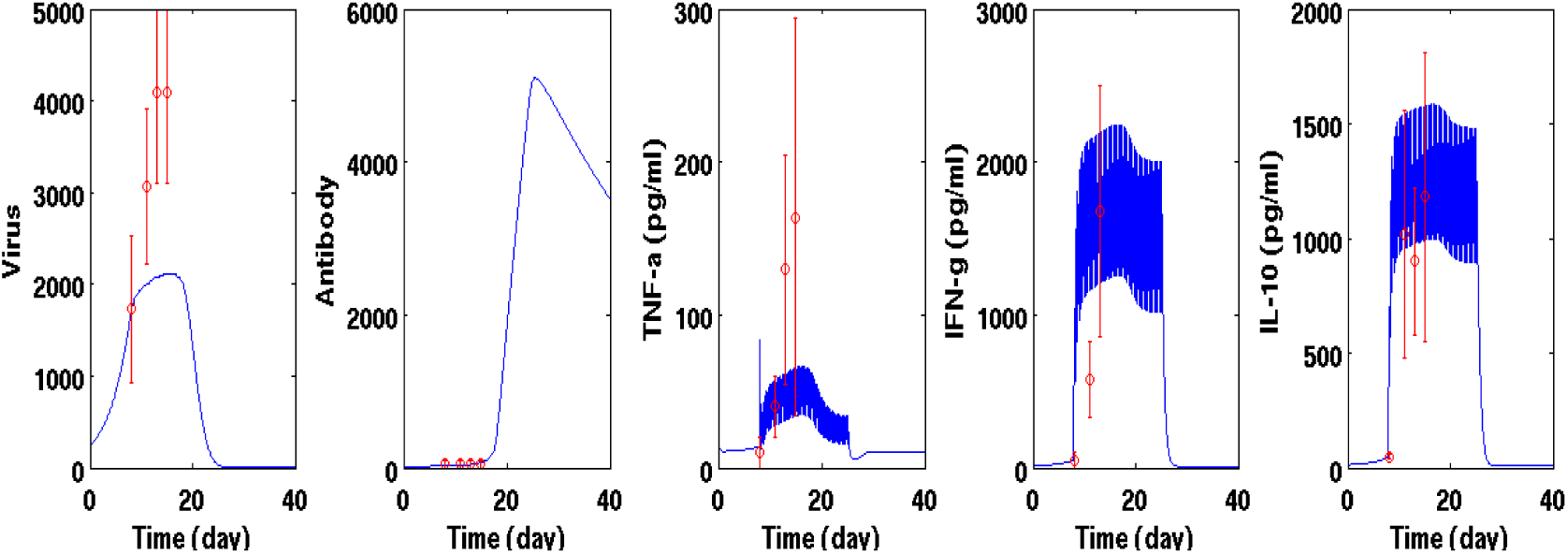
An example of the effects of the immune-interfering therapy. Exogenous IFN-γ (a pulse of 1000 pg/ml every 8 hours) and IL-10 (a pulse of 600 pg/ml every 8 hours) were given to the *in silico* fatal patient to inhibit viral replication and suppress undesirable strong TNF-α upregulation. The *in silico* therapy started from day 8 and ended on day 25. Red circles with error bar are data of the fatalities without the immune-interfering therapy. Model parameter values were taken from the best fit in Fig. 2.

The above results were based on the best fit from H3 and are consistent to those obtained from H5 (Fig. S5). We conclude that a therapy targeting the support and acceleration of T cell functionality should be accompanied with an intricately balanced anti-inflammatory treatment, as a medium level of TNF-α is necessary for antibody induction.

## Discussion

Knowledge of human EBOV infections is limited. Although multiple lines of evidence pointed to the importance of a properly balanced early innate immune response^3, 12, 20, 26^, the requirements for such a proper response in infected humans were not clear. Specifically, it was not clear whether a proper response depends on certain properties of the virus or the host. By calibrating a cytokine network to data from three groups of patients, i.e. fatal cases, survivors, and asymptomatic infections, we have shown here that the differences among the three groups cannot be explained by differential initial viral doses, viral replication rates and viral removal rates. They can be explained, however, by differential immune responses. Specifically, a differential innate immune response alone (H3) explained the data better than differential innate and adaptive immune responses (H5) did. In both H3 and H5, the rate of TNF-α induction by EBOV (*Q*_vf_) in the three groups showed a sequence consistent with disease severity, which was not observed for other parameters. These results suggest that the fate decision of an EBOV infected patient relies on the ability to induce an early innate immune response, which subsequently triggers adaptive responses before it is too late. It is remarkable that despite the fate decisive role of the innate response, virus clearance completely relies on IFN-γ and antibody dependent immune responses.

The reported inoculation period of EBOV infection is in the range of 2-20 days. The inoculation period associated with the data used in the mathematical model was estimated to be about 7 days^3, 20, 23^. The results still hold true, if we start from an inoculation period of 14 days. However, with 3 days the model failed to fit the data. This delineates the range of validity of the proposed stratification and therapy strategy.

The presented model included a positive auto-feedback of IFN-γ representing the stimulating influence of IFN-γ on the differentiation of naïve CD4+ T cells into Th1 cells and on the development of CD8+ T cells, both of which are the main producers of IFN-γ. While there is evidence that TNF-α induces an autocrine loop stabilizing the expression of pro-inflammatory signals including TNF-α itself^27^, we found that the auto-feedback of TNF-α in the model, which represents the whole innate immune response, is not necessary for numerical optimization. While the reason for this asymmetric model structure remains to be clarified, there is *in vitro* evidence that EBOV-induced pro-inflammatory cytokine secretion from human monocytes^6^ is not altered when a TNF-α antagonist is added to the culture, which is in line with the model structure.

We took IFN-γ mainly as a surrogate of the function of cytotoxic T lymphocytes and Th1 cells. This is, in general, not an accurate approximation, as also NK cells secrete IFN-γ upon activation. However, in the special case of Zaire EBOV infection, NK cells have been shown to be depleted much faster than cytotoxic T lymphocyte in nonhuman primates^12, 28^ and, hence, can be neglected for later control of the virus in comparison to cytotoxic T cells.

The numerical optimization studies suggested that IFN-γ and antibody have a similar virus clearance capacity. This quantification of the capacity of cellular and humoral immunity is consistent with experimental results showing that both CD8+ T cells and antibody can convey protection from Ebola virus^13, 29^, that Ebola vaccine provided high protection against disease^30^, and that exogenous IFN-γ protected mice from lethal mouse-adapted EBOV challenge^31^. However, the model suggested that IFN-γ increases two days later in fatal cases than antibody does in survivors (Fig. 5). A delayed induction of IFN-γ was associated with a weak induction of IFN-γ by virus (*Q*_fg_) or TNF-α (*Q*_vg_) in comparison to IFN-γ auto-induction (*Q*_gg_). This might be related to the known impaired function of human dendritic cells *in vivo*, which fail to secrete cytokines in response to EBOV infection, leading to impaired function and anomalous maturation^10, 12^.

The model predicted the dynamics of TNF-α induction to be a main difference among fatal cases, survivors and asymptomatic infections, which suggests a central role of the innate immune response in the early stages and the subsequent earlier transition to humoral responses *in vivo*. This knowledge can be used to stratify EBOV patients at the time of presentation in the clinics, i.e. at the time of onset of symptoms, into potential survivors and fatal cases depending on the measured blood concentration of TNF-α.

The cases classified as fatal may be rescued by an EBOV therapy. In the model it is possible to rescue the in silico patients by a combined IFN-γ and IL-10 therapy. However, this treatment strategy has to be translated into the in vivo case: Due to the necessary reductionist approach in silico, IFN-γ and IL-10 therapy stand for therapies targeting in vivo immune strategies. Therefore, a one-to-one translation of the findings into clinical therapy is limited. However, the use of mono-cytokine therapies in clinical practice is well established, e.g. TNF blockade for the therapy of autoimmunity^32^, or, type I interferon therapy for the treatment of viral infections like hepatitis^33^ and, at least experimentally, IL-10 therapy as anti-inflammatory approach^34^. Direct IFN-γ therapy is also established for prevention of infection in patients with chronic granulomatosis, an innate immunity defect^35^. Interferon therapy has also been discussed and partly tested for the treatment of Ebola^36^, but also other viral infections, like Dengue, parasitosis^37, 38^, and Tuberkulosis^39^. From a mechanistic point of view, interferon-γ therapy might be replaced by type I interferon therapy in the present context, provided it is given early enough. The paradigm that targeting one single mediator will modulate the whole system in a beneficial way was also followed. For example, application of type I interferon strengthens the anti-viral effector function of the immune system^36, 40, 41^ and is reflected in higher levels of the effector cytokine IFN-γ. This means that the clinical practice also reflects a reductionist approach, substituting main modulators for different parts of effector mechanisms. As our model shows, strengthening the anti-inflammatory pathway, e.g. by IL-10 treatment and/or increasing anti-viral mechanisms, e.g. by substituting type I interferons with the right timing could help to increase survival of Ebola infected individuals.

In conclusion, the model suggests that it possible to stratify patients at high risk of fatal infections early on at the time of onset of first symptoms when patients would normally present themselves at the clinics. Further, the model predicts that it is possible to rescue patients classified as fatal EBOV infections by promoting cytotoxic T cell responses, and avoiding undesirable strong inflammatory responses by strengthening anti-inflammatory pathways. The model therefore confirms preclinical and clinical experience and might help to optimize host-directed therapy for Ebola infection.

## Acknowledgement

This work was supported by the Helmholtz Initiative on Personalized Medicine – iMed.

## Supplementary materials

The supplementary material includes the details of the modeling method, the additional hypotheses that have been tested, the Table of parameter values and the captions of the supplementary figures.

### Method

Under consideration of the available data, we took a reductionist approach in modeling immune responses to EBOV infection. Instead of considering different cell types and their corresponding mechanisms in the immune defense, we took their prototype cytokines as surrogates of various aspects of the function of the immune system. Thus, TNF-α was taken as the surrogate of the innate immune response, IFN-γ as that of Th1 and cytotoxic T lymphocytes, IgG as that of the humoral response and IL-10 as the surrogate of immune-regulatory responses.

Patient data obtained during two Ebola outbreaks in Gabon in 1996 from the same virus strain^1, 2^ were used to evaluate different hypotheses for predictive markers of disease progression. Cytokine levels have been measured since clinical admission, until death or convalescence of the patients. Seven out of seventeen patients (survivors and fatal cases) were infected at the same time so that the time lapse between infection and clinical admission was clearly defined for these patients. For the rest of the patients, the same time lapse was assumed. Given the uncertainty in the inoculation time for these patients, we tested different inoculation times in the model. Antigen and antibody levels were reported as titer (dilution). The IFN-γ level in fatalities dropped drastically 1 day before death, which was attributed to massive intravascular apoptosis^1^. Since the current model is not intended for this extreme situation, and since we were interested especially in the early phase responses, we didn’t include the last data point of IFN-γ in fatalities in our studies. The reported IFN-γ protein level in survivors was inconsistent with the trend of mRNA in peripheral blood mononuclear cells from the same patients. The former showed no activation while the latter showed upregulation during the period of viral clearance and clinic recovery^1^. To clarify the possible influence of the inconsistency in IFN-γ protein and mRNA data in survivors, we always carried out two parallel studies, either with or without IFN protein data in survivors. The results were consistent with each other. Certain reported cytokine levels were in the form “< 20pg/ml”. In this instance, we took 10±10 pg/ml instead in the numerical studies.

In addition to the fatalities and survivors, asymptomatic human infections during the two outbreaks have been identified. We also included the data of asymptomatic infection in our studies. However, it appeared that the IgG antibody was measured with a different protocol^3^. In order to avoid any bias from the protocol of data acquisition, we didn’t include the IgG data associated with the asymptomatic infection. For the asymptomatic cases, the infection time was inferred from reported first exposure to infection.

The interactions between all considered cytokines (Fig. 1) are modeled in a phenomenological way, i.e. by sigmoidal functions of the concentrations of the cytokines while neglecting the mediators and the mechanisms underlying each interaction.

### Virus

Viral replication is modeled by logistic growth, with replication rate *p* and carrying capacity *K*. The virus clearance capacities by TNF-α(*F*), IFN-γ (*G*) and antibody (*A*) are denoted by rate constants *R*_f_, *R*_g_ and *R*_a_, respectively. The equation for virus *V* reads

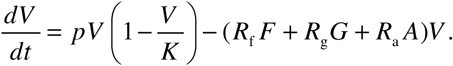

### Cytokines

The dynamics of each cytokine are modeled by a production term and a degradation term. The production term of each cytokine is a Goldbeter-Koshland function (*g*) of the following form:

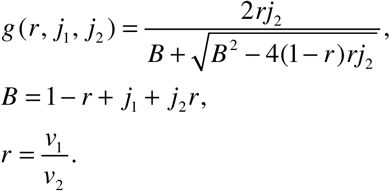

The Goldbeter-Koshland function is a monotonously increasing function of *r*, and has a sigmoidal shape. *r* is the ratio between the effects of activator(s) and inhibitor(s), which are denoted by *v*_1_ and *v*_2_ respectively. In the case of multiple activators (inhibitors), the corresponding term of *v*_1_ (*v*_2_) is composed of linear combinations of each activator (inhibitor), together with the cross-terms up to the second order. The shape of the Goldbeter-Koshland function is determined by *j*_1_ and *j*_2_, which represent the degree of nonlinearity of the activation and inhibition processes, respectively. Note that larger values of *j_1_* (*j_2_*) correspond to lower levels of nonlinearity in the activation (inhibition) process and make the Goldbeter-Koshland function smoother. The equations for the cytokines read:

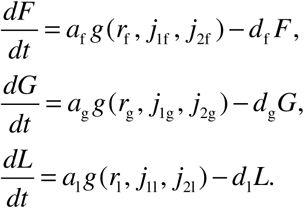

where *F*, *G* and *L* stands for TNF-α, IFN-γ and IL-10, respectively, and

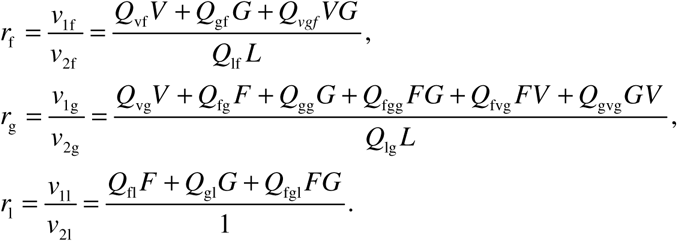

*Q*_xy_ denotes the contribution of x to the induction (inhibition) of y. *Q*_xyz_ denotes the synergistic contribution of x and y to the induction (inhibition) of z and is defined as *Q*_xyz_= *Q*_xz_*Q*_yz_. d_f_, d_g_ and d_l_ are the degradation rates of TNF-α, IFN-γ and IL-10, respectively. *a*_f_, *a*_g_ and *a*_l_ are determined by steady state. The steady state levels of the cytokines were estimated based on the reported values for controls. TNF-α and IL-10 in the controls were reported as “< 20pg/ml” ^24^. IFN-γ in the controls were below the limits of instrument sensitivity ^21^. Consequently, we took 10 pg/ml as the steady state level for each cytokine.

### Antibody

The induction of antibody *A* is induced by TNF-α with a delay. The delay is modeled via the linear train trick (*F*_1_∼*F*_4_). The equations read:

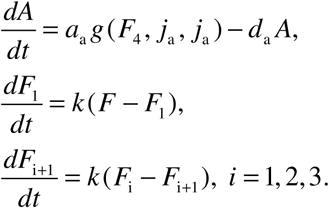

here *d*_a_ denotes the degradation rate of antibody, *k* reflects the delay for antibody induction, which in this case is approximately 4/*k*.

We fitted the cytokine network (Figure 1) to the three data sets (fatalities, survivors, and asymptomatic infections). Specifically, in numerical optimization, we tried to minimize the root mean square (RMS) difference between model simulation (ŷ) and the data (y),

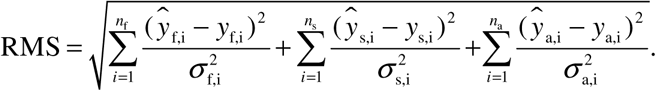

where suffix f, s and a denote fatality, survivor and asymptomatic infection, respectively, σ denotes the standard deviation of the measured data point, and *n* denotes the number of data points.

All the simulation work was done based on the SBPD toolbox for Matlab^4^. The SBPD toolbox was modified in parts for this purpose. A differential evolution based global optimizer was employed to fit the parameters in the model. The optimizer returns a population of parameter-vectors that give rise to acceptable fitting results. Moreover, each optimization task was repeated 100 times. As a phenomenological restriction, we excluded those fitting results where the production rate of any cytokine at resting state was less than one thousandth of the maximal rate. We also varied this restriction to two or five thousand and found no qualitative difference in the results. We took not only the best fitting result for further analyses, but also those fittings that are acceptable based on the Akaike information criterion (AIC) with a relative likelihood more than 0.368 (corresponding to 2 units difference in AIC). We further made sure that the fitting results kept for analysis were well separated in the parameter space in terms of the euclidean distance by filtering out very near solutions.

## Additionally tested hypotheses (related to P5 paragraph 2 in the main text)

A combination of H1 and H2, with the four parameters (initial viral dose, *p*, *R*_f_ and *R*_g_) being different in the three groups, again failed in explaining the data. A variation of H4, with *Q*_vg_ and *j*_1g_ being the differential parameters, failed as well. A variation of H3, where *Q*_vf_ and *j*_1f_ are the differential parameters, explained the data with the relative likelihood of 0.69.

## Supplementary figure captions

**Fig. S1.**
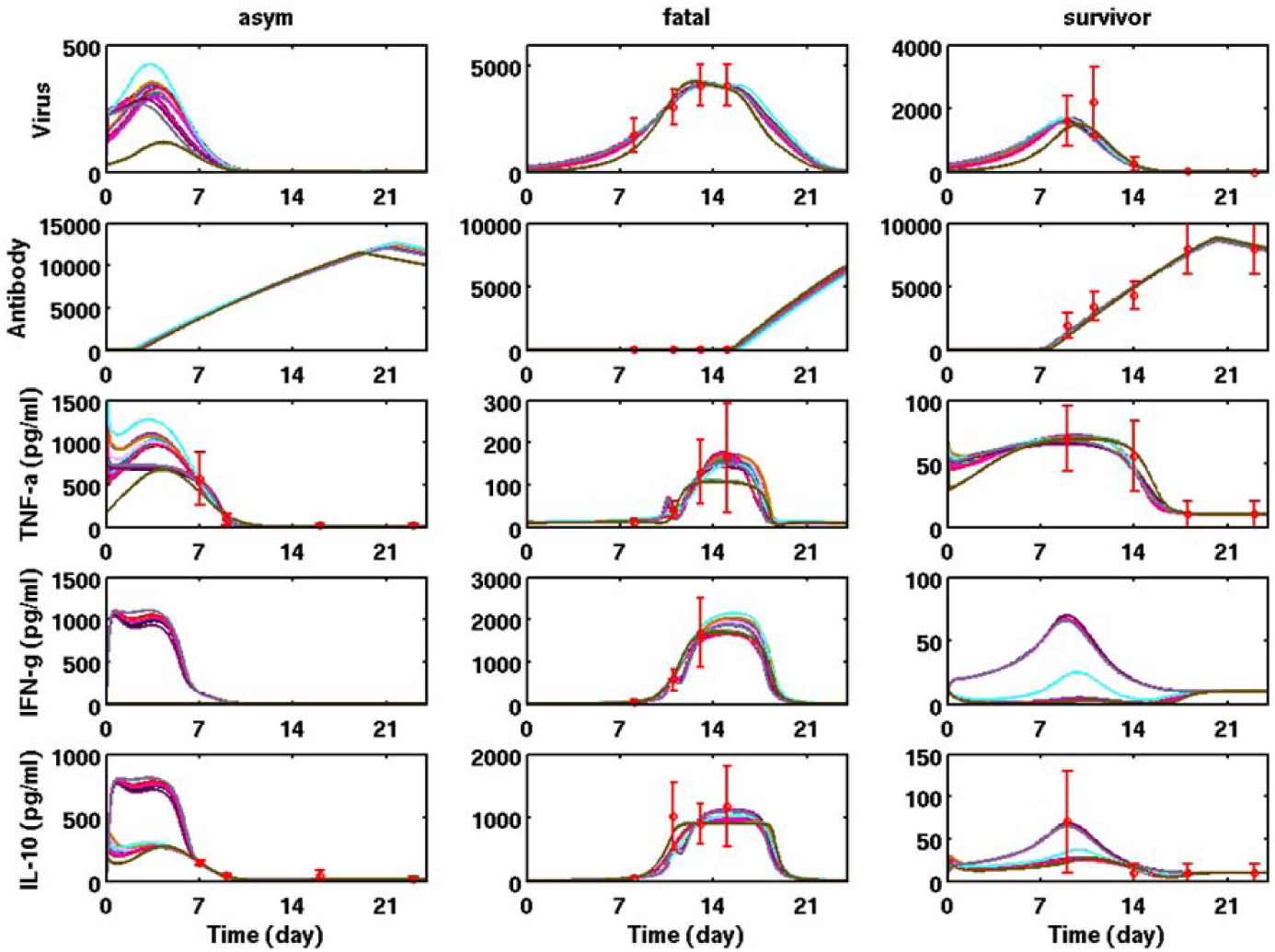
All the accepted fitting results from H3, coded by the color of the curves.

**Fig. S2.**
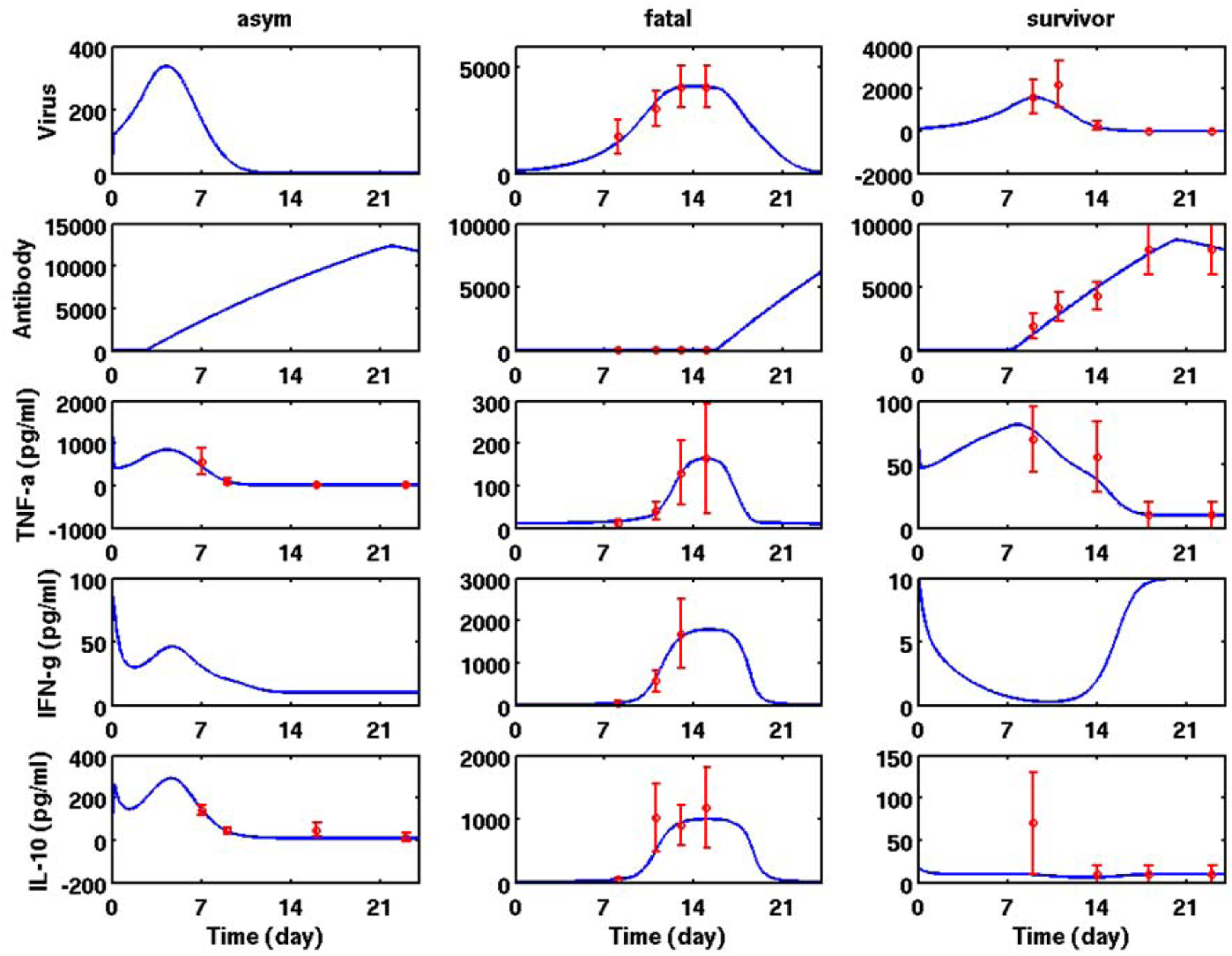
The best fitting results from H5

**Fig. S3.**
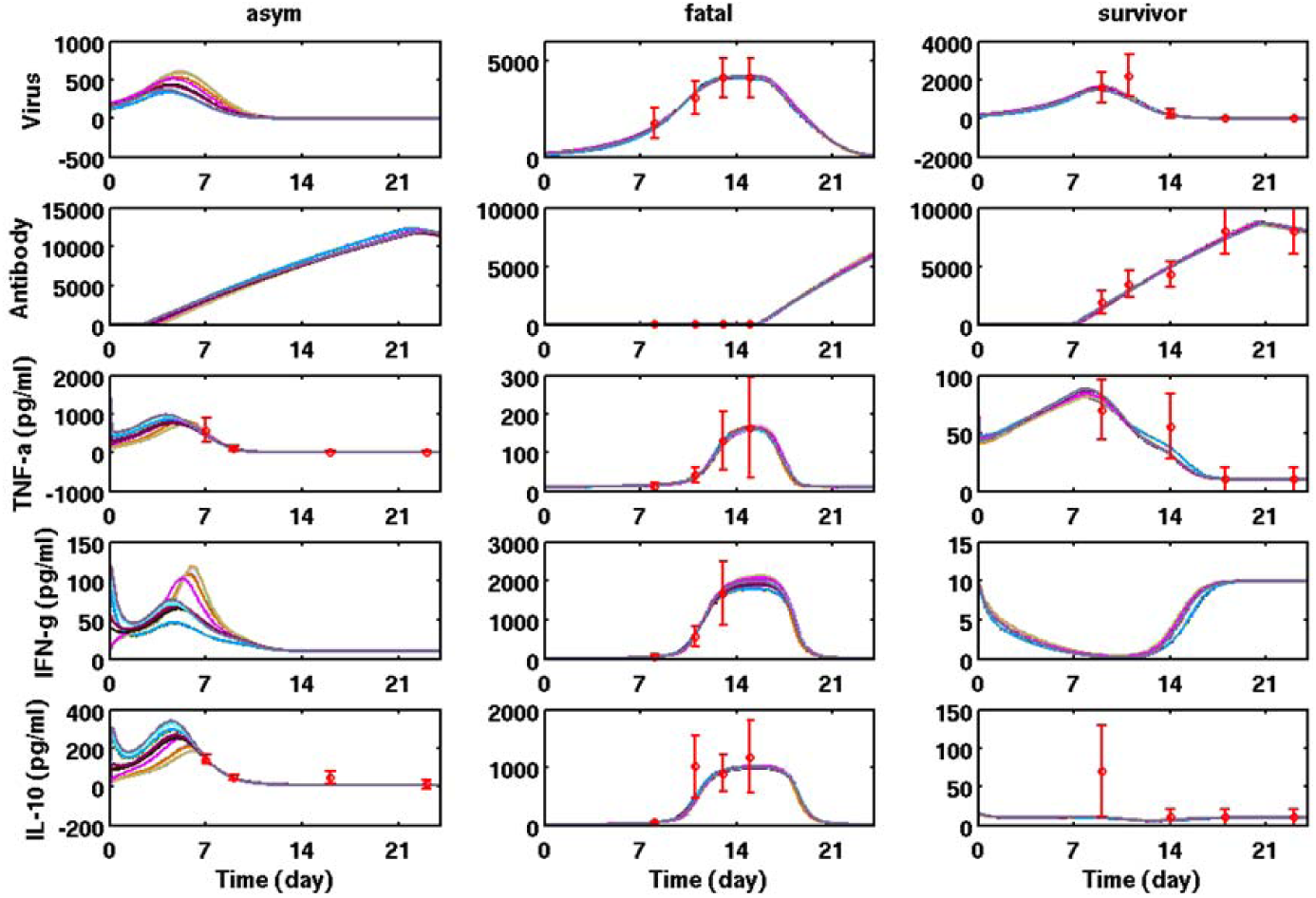
All the accepted fitting results from H5, coded by the color of the curves.

**Fig. S4.**
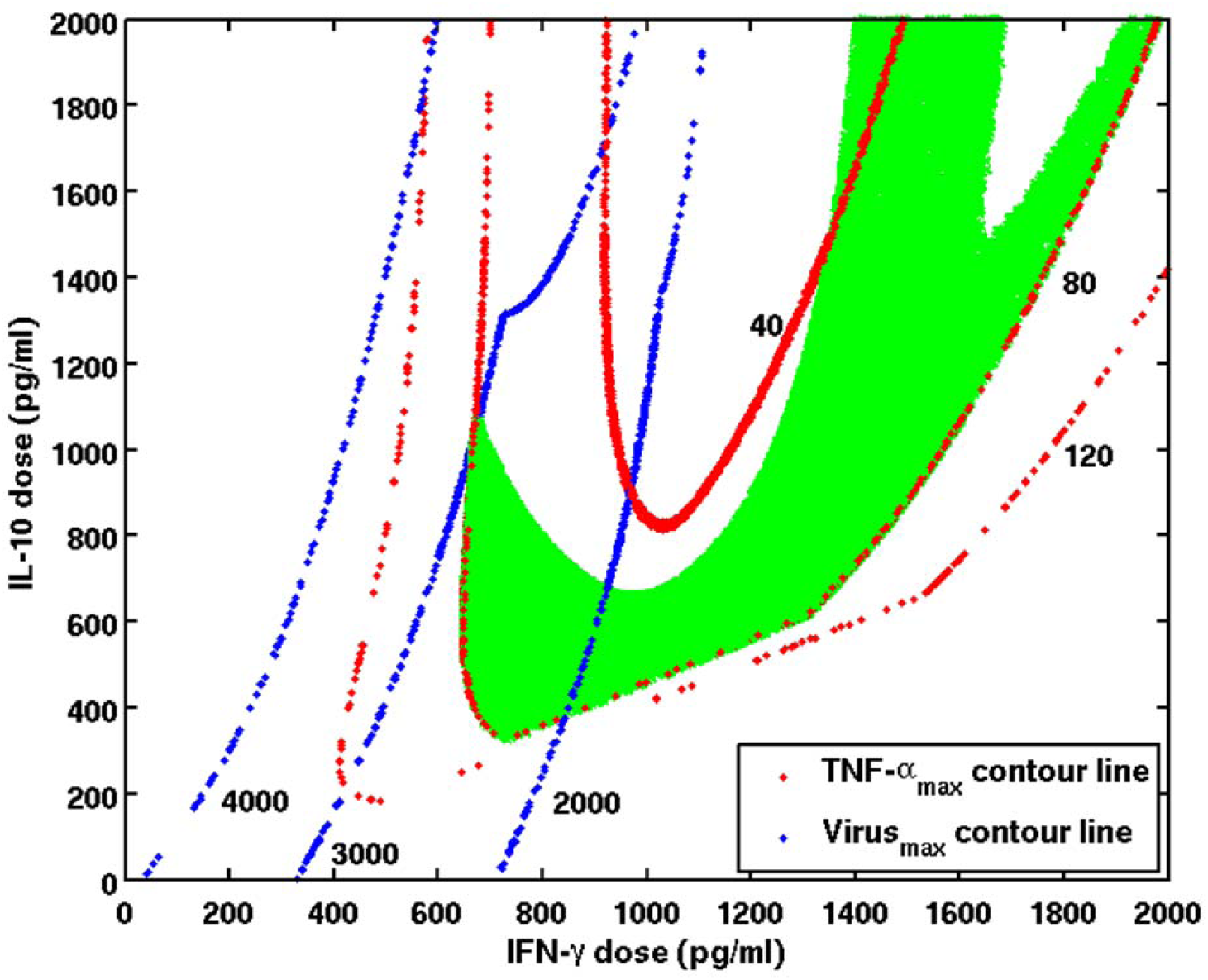
The effect of the immune-interfering therapy depends on the dose of exogenous cytokines. Three contour lines of the maximal viral load (blue) and the maximal TNF-α level (red) during the therapy are plotted. The region is marked green (i.e. successful) if 1) the maximal TNF-α level during the therapy is less than 80 pg/ml; 2) the maximal virus titer is less than 3000; and 3) virus titer is less than 200 when the simulation ended. The therapy started from day 8 and ended when the virus titer became less than 30. The simulations ended on day 43. Model parameter values were taken from the best fit in Fig. 2 in the main text.

**Fig. S5.**
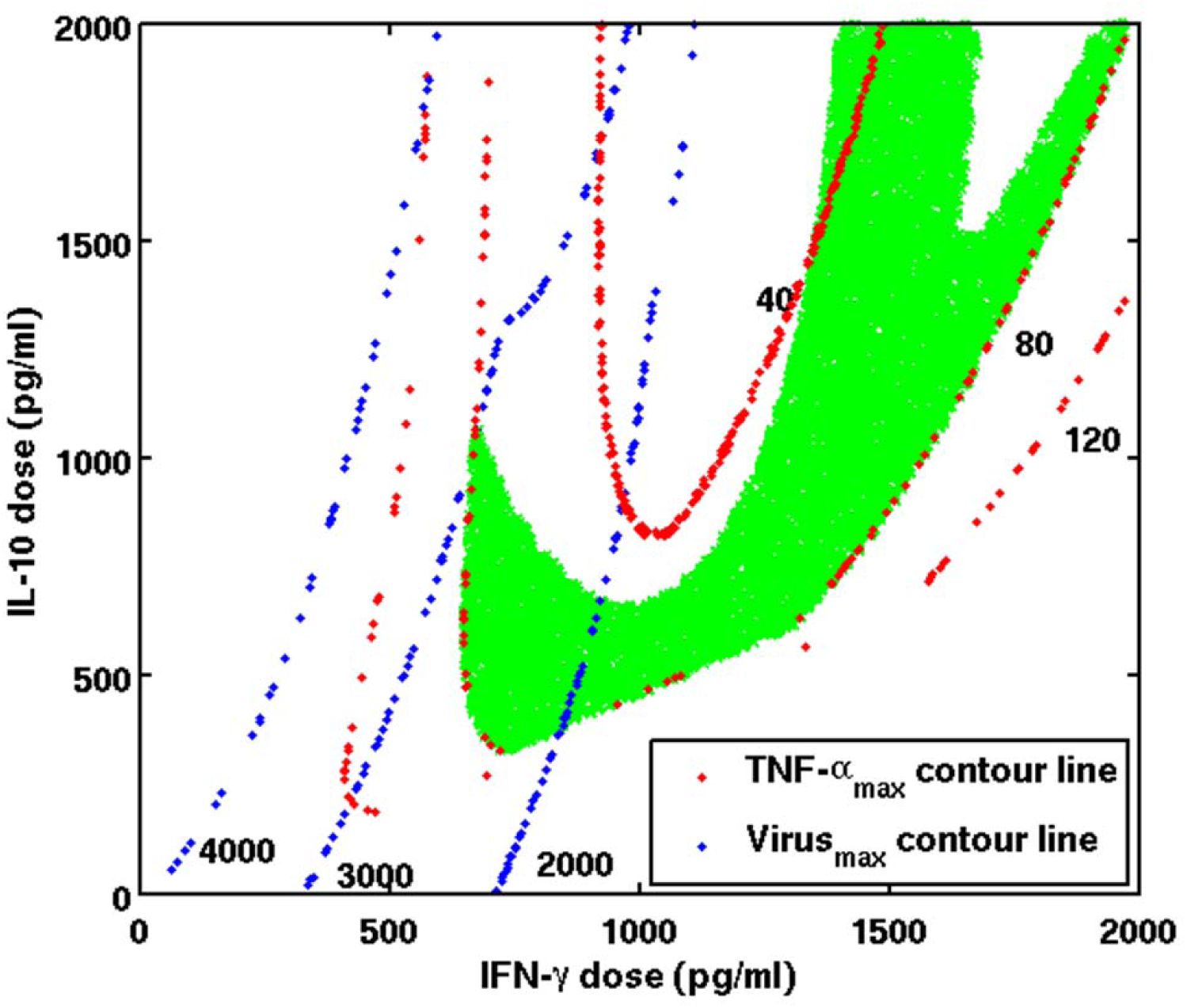
Same as Fig. S4, except that model parameters are based on the best fit from H5.

